# BinDash 2.0: New MinHash Scheme Allows Ultra-fast and Accurate Genome Search and Comparisons

**DOI:** 10.1101/2024.03.13.584875

**Authors:** Jianshu Zhao, Xiaofei Zhao, Jean Pierre-Both, Konstantinos T. Konstantinidis

## Abstract

**Motivation:** Comparing large number of genomes in term of their genomic distance is becoming more and more challenging because there is an increasing number of microbial genomes deposited in public databases. Nowadays, we may need to estimate pairwise distances between millions or even billions of genomes. Few softwares can perform such comparisons efficiently.

**Results:** Here we update the multi-threaded software BinDash by implementing several new MinHash algorithms and computational optimization (e.g. Simple Instruction Multiple Data, SIMD) for ultra-fast and accurate genome search and comparisons at trillion scale. That is, we implemented b-bit one-permutation rolling MinHash with optimal/faster densification with SIMD. Now with BinDash 2, we can perform 0.1 trillion (or ∼10^11) pairs of genome comparisons in about 1.8 hours on a descent computer cluster or several hours on personal laptops, a ∼50% or more improvement over original version. The ANI (average nucleotide identity) estimated by BinDash is well correlated with other accurate but much slower ANI estimators such as FastANI or alignment-based ANI. In line with the findings from comparing 90K genomes (∼10^9 comparisons) via FastANI, the 85% ∼ 95% ANI gap is consistent in our study of ∼10^11 prokaryotic genome comparisons via BinDash2, which indicates fundamental ecological and evolutionary forces keeping species-like unit (e.g., > 95% ANI) together.

**Availability and implementation:** BinDash is released under the Apache 2.0 license at: https://github.com/zhaoxiaofei/bindash

**Supplementary information:** Supplementary data are available at Bioinformatics online.

## Introduction

MinHash, originally developed for detecting duplicate webpages (Broder, et al., 1998), turned out to be a powerful strategy when applied for genome comparisons in the pioneering work from Ondov, et al. (2016): genome kmer set Jaccard index estimated via MinHash can be accurate estimation of genomic distance or ANI via Mash equation ANI or 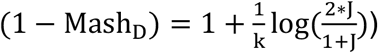. In the original MinHash work, many hash functions are required under the locality sensitivity scheme. An alternative to k-minwise when estimating set similarity with Minwise sketches is bottom-k implementation, where we use one hash function and maintain the sketch as the keys with the k smallest hash values (Broder, 1997). However, it needs a priority queue to maintain the k smallest hash values, and this leads to a non-constant worst-case time per element (overall complexity is O(n*log(k))), which may be a problem in real-time processing of high-volume data streams. More importantly, since only one hash function was used, we are not able to encode set-similarity as an inner product of two sketch vectors. This is because the elements lose their “alignment” – that is, the key with the smallest hash value in one set might have the 10th smallest hash value in another set (Dahlgaard, et al., 2015). Or in other words, the alignment property is preserved if the same components (e.g., value at a given position) of two different sketches are equal with a collision probability that is a monotonic function of some similarity measure, also called a locality sensitive hashing (Shrivastava, 2017). In many real-world applications, such as nearest neighbor search, this property will guarantee theoretically optimal accuracy and search recall will deteriorate significantly if not preserving LSH (Shrivastava, 2017). Another interesting alternative is called One Permutation Hashing, which also applies only one hash function with time complexity O(n + s)). However, it has much larger variance than traditional k-minwise hashing because there might be empty slots in the sketch vector after splitting the sketch into buckets. Densification, that is to fill in the empty slots with some non-empty slots, chosen with certain rules, has greatly improved the accuracy of one permutation hashing and proved to be theoretical equivalent to that of traditional k-minwise hashing. Based on one permutation hashing, original densification (Shrivastava and Li, 2014), improved densification (Shrivastava and Li, 2014) and optimal densification (Shrivastava, 2017) are proposed and have all been proven to be locality sensitive theoretically. We implemented so called original densification and optimal densification in the first version of BinDash (Zhao, 2019). However, more densification strategies have been proposed since the publication of original BinDash, e.g., faster densification (Mai, et al., 2020), re-randomized densification (Li, et al., 2019), bidirectional densification (Jia, et al., 2021) with even better run time behavior. Specifically, faster densification improved the worse-case densification computational complexity in optimal densification from O(n + s^2) to O(n + s*log(s)) with the same average-case O(n + s) (Figure 1b and c) while re-randomized densification further improves accuracy for optimal densification at the cost of additional computation since rerun MinHash within previously empty bins after optimal densification is computationally expensive when there are many empty bins, see detailed complexity analysis for re-randomization densification (Li, et al., 2019). BinDash 2 implemented all flavors and variants of MinHash presented in Broder (1997), Li and König (2010), Li, et al. (2012), Shrivastava and Li (2014), Shrivastava and Li (2014), Shrivastava (2017) (Figure 1a and b) and Mai, et al. (2020) (Figure 1a and c) with SIMD (see below). The implementation detail is presented in Supplementary Material.

**Figure 1.**
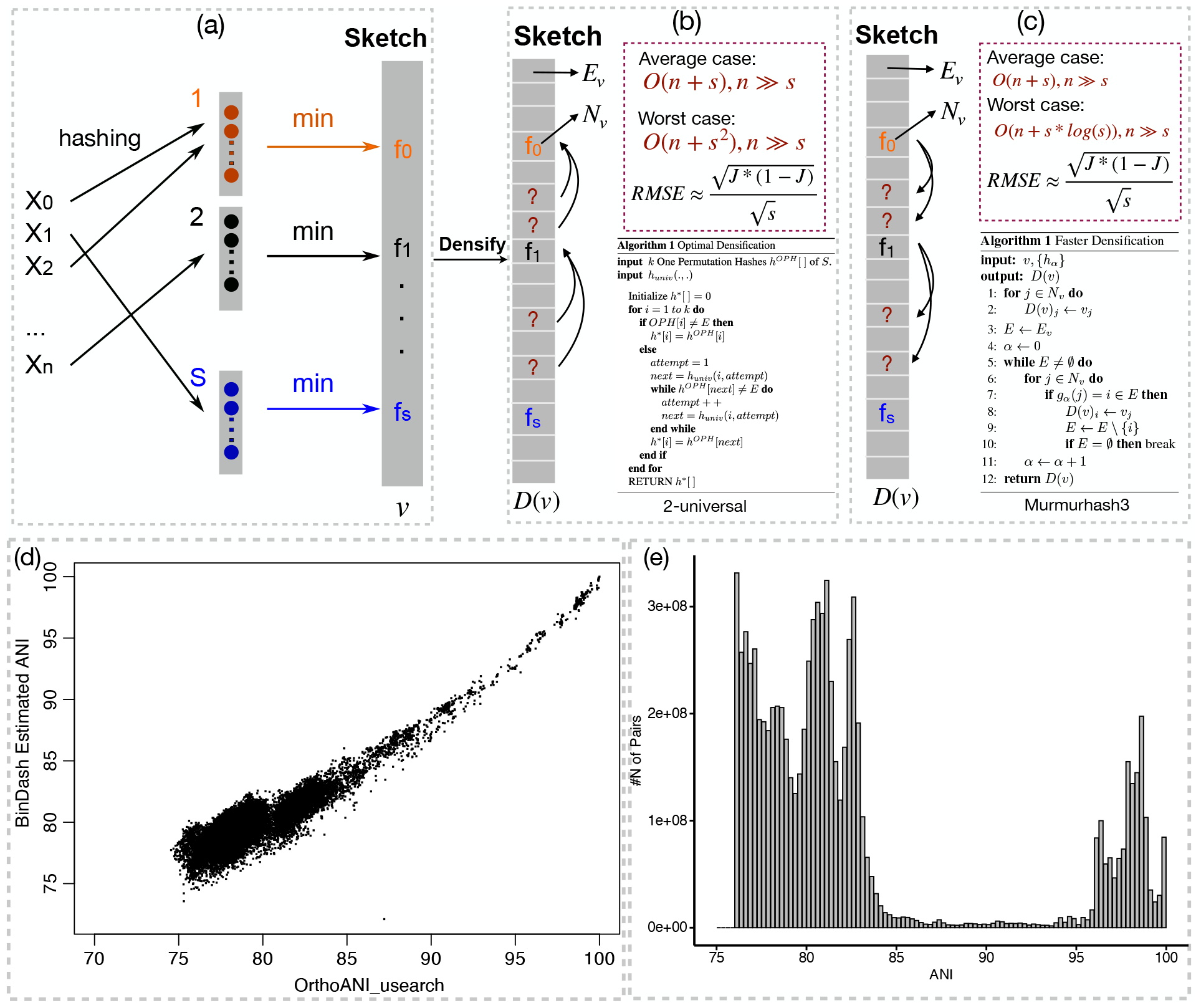
**(a)** One Permutation Hashing; **(b)** Optimal Densification. Mapping empty bins to non-empty bins and copy values from non-empty bins to empty bins, carefully designed 2-universal hashing is required for mapping, implemented in BinDash 1; **(c)** Faster Densification or Reverse Optimal Densification. Mapping non-empty bins to empty bins and copy values to empty bins from non-empty bins, random hashing library can be used, e.g., murmurhash3, implemented in BinDash 2; **(d)** BinDash estimated ANI vs. OrthoANI for randomly selected 3,000 prokaryotic genomes after supervised learning; **(e)** Distribution of ANI pairs with >75% ANI for all vs. all comparisons of ∼318K NCBI/RefSeq prokaryotic genomes. The histogram follows a bimodal distribution with less than 0.01% of pairs fall between ANI 85% and 95%.

The limiting step of BinDash is popcount, which counts the number of non-zero elements in large bit vectors for estimation of collision probability (after XOR operation). However, recent algorithmic advancements in Simple Instruction Multiple Data has provided the opportunity to further speed up popcount for many instruction sets (e.g., AVX2, AVX512 and SVE512) (Langarita, et al., 2023; Muła, et al., 2018). Since we use 64-bit integer type as sketch vector to store hashes from kmers, it possible to use SIMD to count the number of non-zero elements in parallel after rearranging the sketch vector in a way such that each small portion of sketch vector fill into AVX instructions (e.g., 512 bit for AVX512 or SVE512, 256 bit for AVX2) and also take care of the remaining part that does not fit for any size of sketch vectors.

Genomic distance, measured via Jaccard index can be transformed into genome average nucleotide identity via the Mash equation mentioned above. However, it has been recently proved that the Poisson model assumption of sequence evolution can also be replaced by Binomial model, which give more accurate estimation of ANI for distantly related genomes (e.g., below 85% ANI) (Belbasi, et al., 2022): 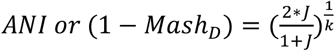. We also add this Binomial model option in BinDash2. To further improve the accuracy of BinDash 2 estimated ANI when compared to alignment-based ANI, we implemented a supervised leaning step, which minimize the RMSE between BinDash 2 estimated ANI according to the RMSE equation: 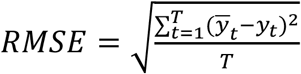 where 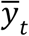 and *y*_*t*_ are BinDash 2 estimated ANI and orthoANI (usearch) ANI for each pair, respectively. T is the total number of pairs of genomes in the training dataset. We use a large collection of training genomes (10,000) extracted from NCBI/RefSeq, covering ANI from 75% to 100%.

Similar to the first version, for each set, BinDash2 first applies one-permutation MinHash (Li, et al., 2012), which use one predefined hash function to all elements/kmers in the set. Then, one-permutation MinHash deterministically partitions the hash value universe into a predefined maximum number B of buckets, extracts the smallest hash value in each bucket, and extracts the b lowest bits (b=14 in practise) of each smallest hash value (Li and König, 2010). These B*b bits are use as the signature of the set. Usually, a hash value v is assigned to the ⌈v/(M/B)⌉ bucket, where M is the maximum possible value for v. One-permutation MinHash may produce a bucket that contains hash values for one set but no hash values for another set, we then apply densification algorithms (Shrivastava and Li, 2014). In additional to all MinHash schemes implemented in original version (original MinHash, bottom-k MinHash and optimal densification), we then implemented faster densification in BinDash2. That is, we now implemented b-bit one-permutation rolling MinHash with optimal/faster densification with SIMD. Specifically, cyclic-polynomial rolling hash based on iterated string hashing is much faster than MurMurHash3 for DNA strings as in Mash (Lemire, 2012; Lemire and Kaser, 2014). We also added nearest neighbor search option to only report nearest genomic distance to query genomes with the consideration that the three densification schemes are all LSH, a property important for genome search and classifications, questionable for other similar softwares without LSH properties such as Mash, Dashing and Sourmash (Baker and Langmead, 2019; Brown and Irber, 2016; Ondov, et al., 2016).

## Evaluation

We compared BinDash2 with BinDash, Mash, Dashing 1 and 2, the state-of-the-art MinHash-based bioinformatics softwares. On Dec. 22, 2023, the 315,686 assemblies of bacterial and archaea genomes in RefSeq were downloaded. The downloaded genomes consist of 412.7 GB of gzip-compressed FASTA files (∼2.1 terabytes for raw fasta files). The 315,686 compressed genomes are used as input data to each software. For each software, we recorded the following: total size of the files used to represent sketches and wall-clock runtime of each command. Each software ran, with its default parameter values (other than a cutoff of 0.2 for mutation rate) with 24 threads, on a Red Hat Enterprise Linux (Intel(R) Xeon(R) Gold 6226 CPU @ 2.70GHz, supports AVX2 and AVX512 instruction sets). BinDash2, dashing 1 and 2 and Mash are both composed of two commands: sketch and dist. The command sketch compute sketches based on hashing kmers from genomes. The command dist compares each sketch used as query to each sketch used as target. The total runtime of these two commands is the total runtime of the corresponding software. In all comparisons, we use sketch size 10,000 instead of the default 2000 to have accuracy at 99% ANI or above, sketch size is important for real-world genomic distance comparisons for species level ANI comparisons (e.g., 95% or 99%) as shown in many places (Jain, et al., 2018). For 318,756 bacterial and archaea genomes, BinDash 2 is 47.8% faster than original BinDash, ∼80 time faster than Mash, ∼8 and ∼17 times faster than Dashing 1 (HLL) and 2 (ProbMinHash or SetSketch) respectively, for the dist command (Table S1) while for sketch, BinDash 2 is sighly slower than Dashing 1 (HLL, the fastest). However, since sketch step is always less than 20 minutes for ∼318K genomes, it is not a limiting step for all tools mentioned above (Table S2). In terms of sketch size stored on disk, BinDash 2 and BinDash is about 3 times smaller than Mash but larger than Dashing (Table S1).

We did a theoretical analysis of RMSE for all MinHash-like and HyperLogLog-like algorithms and showed that MinHash-like algorithms are generally more accurate than HyperLogLog-like algorithms in terms of estimating Jaccard index (Table S2). We computed the true pairwise Jaccard indices of the 120 reference genomes chosen among these 318,756 genomes with ANI above 80%. The true Jaccard indices serve as ground truth and root-mean-square error (RMSE) was used to measure the accuracy of all tools. BinDash 2 remains almost the same accuracy with original BinDash, with RMSE better than both Mash and Dashing (Table S3). More importantly, BinDash 2 and Mash RMSE converges to 0 as sketch size increases in theory while for Dashing 1 and 2 (HyperLogLog), there is no such guarantee (Table S2) (Flajolet, et al., 2007; Gakhov, 2022).

To show the real-world application of BinDash 2, we first compare the ANI from BinDash 2 with orthoANI after correction: A correcting factor σ is obtained via the supervised learning step to correct the final ANI or 1-Mash_D_ so that the final output ANI value correlates well with orthoANI (usearch) ANI (Figure 1d). Then we applied it to defining bacterial genome species boundaries by performing all versus all comparisons among ∼318K genomes. We see a clear 85% to 95% ANI gap (Figure 1e), consistent with more accurate but much slower software called fastANI. We believe this consistent gap is not sampling bias or cultivation bias because many genomes deposited recently are environmental genomes obtained by metagenomics and they reject cultivation and isolation.

## Discussion

BinDash 2 implemented the fastest densification idea called faster densification, which has the same theoretical RMSE with traditional MinHash. Overall, MinHash-like algorithms for estimation of Jaccard index are more accurate than HyperLogLog-like algorithms as implemented in Dashing 1 and Dashing 2. To further improve the accuracy of MinHash estimated Jaccard index or ANI, it is possible to explore new MinHash algorithms like re-randomized MinHash (Li, et al., 2019) and circulant MinHash (Li and Li, 2022), which are all theoretical breakthroughs very recently. However, both algorithms achieved smaller RMSE (using same sketch size) at the expense of additional computations. For example, re-randomized MinHash requires additional MinHash step within each empty bin after running optimal densification, which can be several times slower than faster densification according to theoretical analysis (Li, et al., 2019). Circulant MinHash requires large number of random permutations for the second permutation procedure, which is not so efficient for large number of bins in sketch vector in practice. However, we can use slightly larger sketch size in fast densification to achieve similar accuracy with re-randomized MinHash or Circulant MinHash because the running time of faster densification is not compromised due to the fact that average-case O(n + s) is not affected by sketch size s (n is always more than 100 times larger than s for genomic applications). In this regard, BinDash 2 achieved the best running time and accuracy trade-off among all MinHas-like algorithms, including newly invented ones.

We have also showed that BinDash 2 can be used to perform large scale genome comparisons and help define prokaryotic genome species. Taken together, via implementing new algorithms and computational optimization, we believe BinDash 2 will be a practical alternative to similar tools and will help very large-scale microbial genome search and comparisons while maintaining accuracy for biological knowledge discovery.

